# Optical resolution is not the limiting factor for spatial precision of two-photon optogenetic photostimulation

**DOI:** 10.1101/2023.07.01.547318

**Authors:** Robert M. Lees, Bruno Pichler, Adam M. Packer

## Abstract

Two-photon optogenetics combines nonlinear excitation with noninvasive activation of neurons to enable manipulation of neural circuits with a high degree of spatial precision. Combined with two-photon population calcium imaging, these approaches comprise a flexible platform for all-optical interrogation of neural circuits. However, a multitude of optical and biological factors dictate the exact precision of this approach *in vivo*, where it is most usefully applied. Here, we carefully assessed how the optical factors determine the spatial precision of activation. We found that optical resolution is not the limiting factor of the spatial precision of two-photon optogenetic photostimulation, and by doing so, reveal the key factors to improve to achieve maximal precision. Our results enable future work to focus on the optimal factors by providing key insight from controlled experiments in a manner not previously reported. This research can be applied to advance the state-of-the-art of all-optical interrogation, extending the toolkit for neuroscience research to achieve spatiotemporal precision at the crucial levels at which neural circuits operate.

## Introduction

Interrogating neural circuits requires observation and intervention at the spatiotemporal scales that these circuits operate, i.e. spatially at the cellular level (∼10 µm) and temporally within milliseconds. Two-photon optogenetics^1–6^ refers to the use of two-photon excitation to photostimulate opsin proteins embedded in the cellular membranes in order to activate or inhibit cells in a targeted fashion. The use of two-photon excitation enables targeting at the scale of individual neurons deep in scattering tissue, for example, up to ∼0.5 mm deep in the brains of awake, behaving rodents. This approach extends the optogenetic toolkit beyond bulk activation using fibre-coupled LEDs by enabling researchers to target cells with greater specificity. For example, neurons can be targeted based on their functional properties instead of, or in addition to, genetic or viral targeting approaches. Combined with holography for multiplexed targeting, and calcium imaging for neural activity recording, two-photon optogenetics becomes a versatile all-optical interrogation^7–9^ approach for reading and writing neural activity *in vivo* with unprecedented precision.

Since the beginning of two-photon optogenetics, a key goal has been to target neurons for activation with single-cell precision. One of the key metrics for quantifying single-cell precision is the physiological point spread function (PPSF), which was defined by Pégard et al 2017^10^ as “the photocurrent response to multiphoton photostimulation as a function of the displacement between the holographic target and the patched cell”. First, a recording of neural activity from a single neuron is established either by patch-clamp electrophysiology, the gold standard for recording subthreshold depolarisation and/or spiking activity, or by calcium imaging, a widely used indirect recording technique for spiking activity. Next, the photostimulation light is directed to the neuron to elicit a measurable response (i.e. photocurrent, action potential, or calcium transient). Finally, the light is moved away from the cell in a stepwise fashion laterally and axially to determine the neural responses as a function of distance between the light and the targeted neuron. One key summary metric from such recordings is the full width at half max (FWHM) of the response versus the axial dimension, because the native resolution of two-photon excitation produced at the focus of an imaging system is generally worse in the axial than the lateral dimension. The best case scenario would be that the axial FWHM of the PPSF is approximately equal to the diameter of the neuron being targeted, such that if the light is half of one cell’s diameter away from that cell, the activation of that neuron drops to half of its maximal value (assuming that opsin is spread uniformly around the cell body). This optimum may not be achieved if the cell is not a perfect sphere, which cells rarely are when in a native tissue environment.

The factors that affect the PPSF can be separated into three categories: optical, optogenetic, and cellular. The optical factors are: the properties of the optical point spread function (OPSF) that delivers light to the sample, how much laser power is directed at the sample, the laser exposure time, and whether the beam is scanned across the cell in a specific way such as a raster or spiral. The optogenetic factors are: the number of opsin proteins activated, the conductance of each, and where they are located in the cell. The cellular factors are: the size and shape of the cell, and the depolarisation required to reach action potential threshold. The action potential threshold depends on the cell’s intrinsic electrophysiological properties such as resting membrane potential, input resistance, and ion channel expression. There are, of course, other factors that affect such experiments such as the wavelength of light used, the number of targets that can be simultaneously addressed, and so on – but these factors do not directly affect the PPSF beyond the key factors described above.

Researchers have attempted to optimise each of these categories of factors. Regarding the optical factors, researchers have employed advanced light shaping approaches such as temporal focusing^11, 12^. Regarding the optogenetics factors, researchers have developed stronger opsins^13–15^ and employed targeted expression strategies such as perisomatic restriction^16–18^. Regarding the cellular factors, researchers have tailored the activation to each cell^19–21^ to optimally reach that cell’s threshold.

A large number of papers have reported both optical PSF and physiological PSF results under a wide range of optical and biological conditions (see Table 1). In general, the PPSFs are larger than the OPSFs, and also larger than the desired outcome, i.e. they are larger than the size of the neurons being targeted. Mouse neurons are on the order of 10-20 µm in diameter, but the PPSFs are generally in the 20-40 µm range, with recent papers employing perisomatic restriction *in vivo* all hovering around 30 µm despite using different optical targeting approaches and opsins. Zebrafish neurons are smaller, generally less than 10 µm, but again the reported PPSFs from papers targeting these neurons are 10-15 µm.

**Table 1:** Physiological point spread functions (PPSFs) from papers^2, 3, 5–8, 10, 13, 14, 18, 22–32^ using two-photon optogenetic excitation of neurons. The lowest axial FWHM values tend to come from papers using perisomatic restriction and/or temporal focusing. Note the best consistent results from zebrafish neurons at 10 and 15 µm.

| <u>Paper</u> | <u>Year</u> | <u>Slice</u><br><u>or</u><br><u>in vivo</u> | <u>Type of cell</u> | <u>Opsin</u> | <u>Perisomatic</u><br><u>restriction?</u> | <u>Measurement</u> | <u>Method</u> | <u>Axial</u><br><u>FWHM</u><br><u>($\mu\text{m}$)</u> |
| --- | --- | --- | --- | --- | --- | --- | --- | --- |
| **Rickgauer** | 2009 | Slice | HEK293T | ChR2 | No | Photocurrent | Spiral | ~30 |
| **Papagiakoumou** | 2010 | Slice | Mouse pyr. | ChR2 | No | Photocurrent | TEFO | 26 |
| **Andrasfalvy** | 2010 | Slice | Rat & mouse pyr. | ChR2 | No | Photocurrent | TEFO | ~30 |
| **Packer** | 2012 | Slice | Mouse pyr. | C1V1(T) | No | Spiking | Raster | 30 |
| **Prakash** | 2012 | Slice | Mouse pyr. | C1V1(T) | No | Spiking | Raster | 13 |
| **Rickgauer** | 2014 | In vivo | Mouse CA1 | C1V1(TT) | No | Optical | TEFO | 20 |
| **Packer** | 2015 | In vivo | Mouse pyr. | C1V1(T) | No | Spiking | Spiral | 47 |
| **Baker** | 2016 | Slice | Mouse pyr. | ChR2-Kv2.1 | Yes | Spiking | TEFO | 23 |
| **Carrillo-Reid** | 2016 | In vivo | Mouse pyr. | C1V1 | No | Optical | Spiral | ~30 |
| **Dal Maschio** | 2017 | In vivo | Zebrafish | ChR2 | No | Optical | CGH | 10 |
| **Shemesh** | 2017 | Slice | Mouse V1 | soCoChR | Yes | Photocurrent | CGH | 43 |
| **Shemesh** | 2017 | Slice | Mouse V1 | soCoChR | Yes | Photocurrent | TEFO | 24 |
| **Pegard** | 2017 | Slice | Mouse pyr. | ST-Chronos or<br>ST-ChrimsonR | Yes | Spiking | 3D-SHOT | 28 |
| **Pegard** | 2017 | In vivo | Mouse pyr. | ST-Chronos or<br>ST-ChrimsonR | Yes | Spiking | 3D-SHOT | 29 |
| **Pegard** | 2017 | In vivo | Mouse pyr. | ST-ChrimsonR | Yes | Spiking | 3D-SHOT | 29 |
| **Mardinly** | 2018 | In vivo | Mouse pyr. | ST-ChroME | Yes | Spiking | 3D-SHOT | 28 |
| **Forli** | 2018 | In vivo | Mouse pyr. | ChR2-Kv2.1 | Yes | Spiking | CGH | 34 |
| **Yang** | 2018 | In vivo | Mouse pyr. | C1V1 | No | Spiking | Spiral | ~40 |
| **Yang** | 2018 | In vivo | Mouse pyr. | C1V1 | No | Optical | Spiral | ~40 |
| **Marshel** | 2019 | In vivo | Mouse pyr. | ChRmine-Kv2.1 | Yes | Optical | Spiral | 26 |
| **Marshel** | 2019 | In vivo | Mouse pyr. | ChRmine-Kv2.1 | Yes | Optical | Spiral | 28 |
| **Gill** | 2020 | In vivo | Mouse mitral &<br>granule | ChrimsonR | No | Optical | CGH | 60 |
| **Dagleish** | 2020 | In vivo | Mouse pyr. | C1V1-Kv2.1 | Yes | Optical | Spiral | 40 |
| **McRaven** | 2020 | In vivo | Zebrafish | CoChR-Kv2.1 | Yes | Spiking | CGH | 15 |
| **Daie** | 2021 | In vivo | Mouse pyr. | ST-ChrimsonR | Yes | Optical | Spiral | 80 |
| **Forli** | 2021 | In vivo | Mouse pyr. | stCoChR | Yes | Spiking | CGH | 42 |
| **Bounds** | 2021 | Slice | Mouse pyr. | ST-ChroME | Yes | Spiking | 3D-SHOT | 28 |
| **Oldenburg** | 2022 | In vivo | Mouse pyr. | ST-ChroME or<br>ChroME2s | Yes | Optical | 3D-SHOT | 30 |
| **Jin** | 2022 | In vivo | Mouse pyr. | ChRmine-Kv2.1 | Yes | Optical | CGH<br>(beaded<br>ring) | 26 |

It is unclear which factors can be further optimised to match the PPSF to the approximate size of the neuron being targeted. It is also unclear which factors are already nearly optimal under the variety of experimental conditions used. We sought to answer one specific aspect of these general queries: how much optical resolution is necessary? We filled this gap in knowledge by performing a controlled experiment varying only the optical resolution while holding all other factors constant. We found there is no effect when halving the FWHM of the OPSF (from 19 to 8 µm). We hypothesize that this is primarily because the cell size and shape determine the PPSF under these conditions. Additionally, we are likely working in a condition where out-of-focus excitation is occurring.

## Results

### Varying the photostimulation OPSF axial spread using a spatial light modulator

We used a spatial light modulator (SLM) to vary the size of the photostimulation OPSF and control its z-position relative to the targeted neuron. To elongate the point spread function, we cropped the phase masks being applied to the SLM active area (**Figure 1a**), effectively underfilling the back focal plane of the objective. The phase masks produced a single spot, roughly 100 µm from the centre of the photostimulation field-of-view. We imaged sub-micron fluorescent beads using the 1030 nm photostimulation beam in the SLM path to measure the OPSF of the excitation produced by these phase masks. In the axial dimension, the ‘small’ OPSF had a full-width at half-maximum (FWHM) of 8.3 µm and the large OPSF had a FWHM of 18.6 µm (large OPSF, n=3 beads; small OPSF, n=4 beads; **Figure 1b**). We varied the z-position of the OPSF in 15 µm steps from -60 µm to +60 µm relative to the imaging plane by creating new phase masks for each position. This was confirmed by burning the resultant holograms into a plastic slide and measuring the z-position of the burns relative to the imaging plane using the motorised stage (data not shown). In summary, we created a large and small OPSF that we could position in three dimensions to target individual neurons and test the effect of OPSF size on photostimulation response.

**Figure 1:**
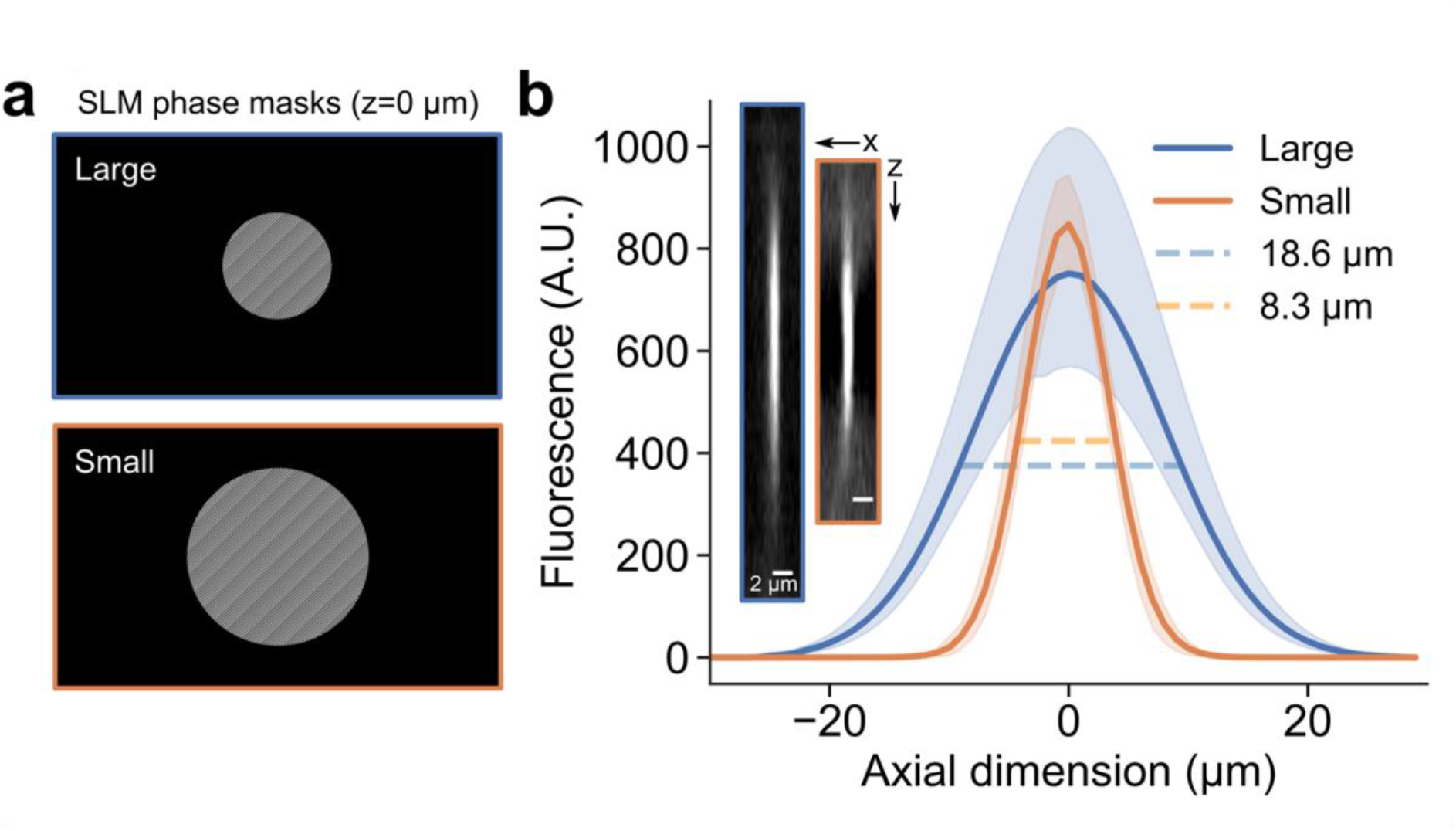
Varying the axial spread of the photostimulation OPSF using a spatial light modulator. **(a)** Example phase masks applied to the SLM active area. These phase masks were used to vary the axial spread of OPSFs, the examples are from the 0 µm photostimulation z-position. **(b)** Mean Gaussian curves +/- 95% C.I. (confidence interval) fitted to the z-profile of 0.2 µm fluorescent beads imaged using the photostimulation OPSF (large OPSF, n=3 beads; small OPSF, n=4 beads). The full-width at half-maximum is indicated (*dashed line*), showing the axial extent of the OPSF. *Inset*, example images from a bead imaged with the ‘large’ (*blue outline*) and ‘small’ (*orange outline*) OPSFs. The images are xz projections; brightness and contrast have been adjusted separately for the two images to highlight the shape.

### Optimising photostimulation for detection of non-saturated neuronal responses

We needed to detect small changes in a neuron’s response to photostimulation to produce a curve of neuronal response versus z-position. To do this, we optimised the average power and frequency of photostimulation to prevent saturation of the neuronal response (as measured using calcium indicators). In this study, we used the normalised change in fluorescence (ΔF/F) of a common calcium indicator, GCaMP6s, as a proxy for neuronal response. We expressed GCaMP6s in neurons in the somatosensory cortex of mice alongside a peri-somatic red-shifted opsin, C1V1-Kv2.1, conjugated to the fluorescent protein mScarlet (**Figure 2a**). We identified single mScarlet-positive neurons in the field of view to target for photostimulation.

**Figure 2:**
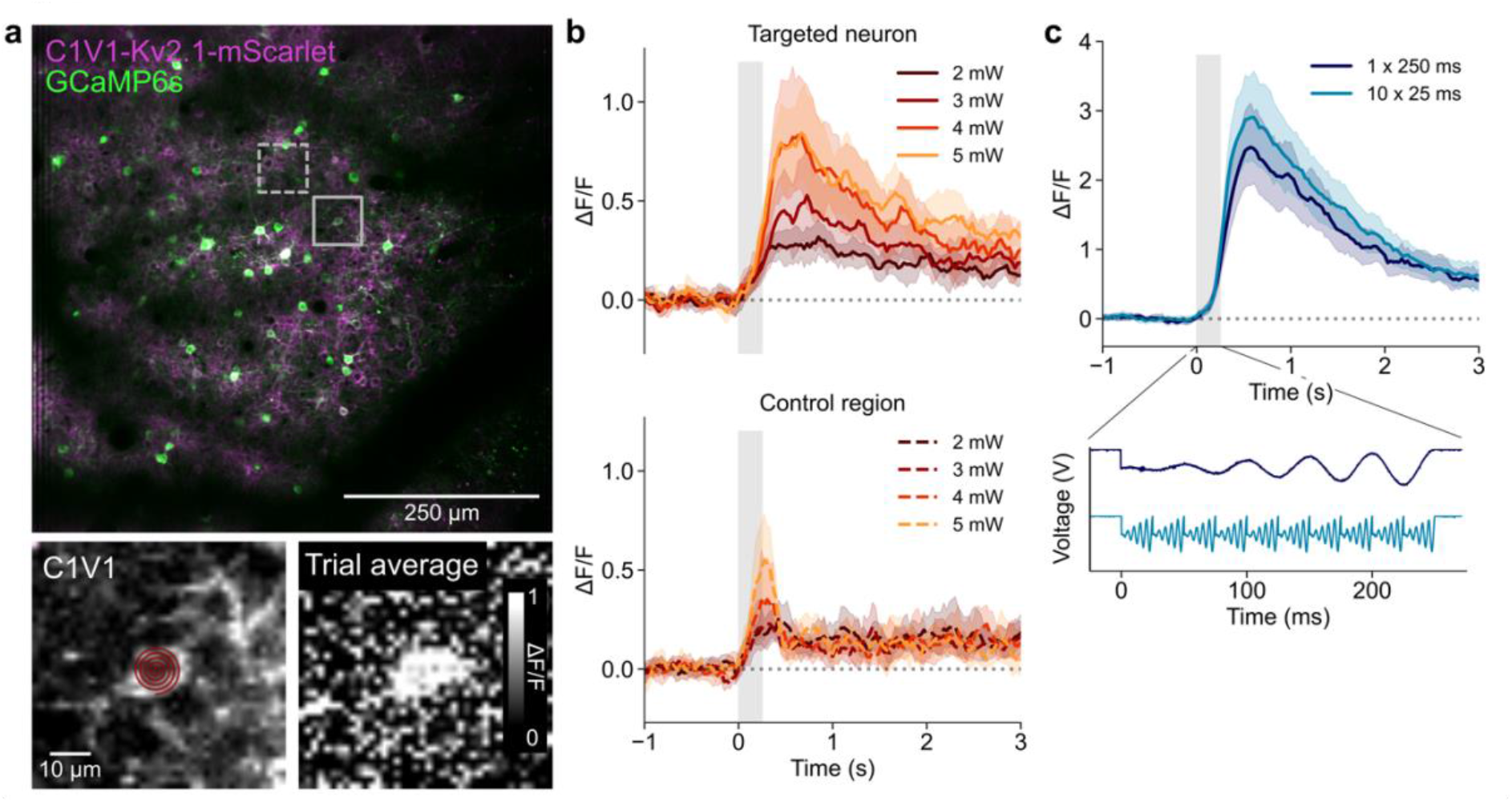
Optimising photostimulation parameters for detection of non-saturated neuronal responses. **(a)** *Top*, example two-photon fluorescent image in layer 2 of the mouse somatosensory cortex. Neurons are labelled through viral expression with GCaMP6s and C1V1-Kv2.1-mScarlet (*solid box*, targeted neuron; *dashed line*, control region). *Bottom-left*, zoom of the targeted neuron showing the C1V1-Kv2.1-mScarlet labelling. An example spiral of 10 µm diameter is overlaid on the targeted neuron. *Bottom-right*, trial-averaged image of the targeted neuron in units of normalised change in fluorescence (ΔF/F) averaged across a period of 1.5 seconds post-photostimulation. **(b)** *Top,* mean ΔF/F traces +/- 95% C.I. (confidence intervals) for all neurons stimulated at each power level (different neurons for each power level; 2 mW, 3 mW, n=10 neurons each; 4mW, 5mW, n=9 neurons each). *Bottom*, mean ΔF/F traces +/- 95% C.I. for all control regions (*dashed box* in panel a). **(c)** Mean ΔF/F traces +/- 95% C.I. for all neurons in each spiral condition (every cell was tested with both spiral conditions, n=24 cells). *Bottom*, voltage traces from the x galvanometric mirror that spirals the photostimulation beam showing the single slow spiral (1 x 250 ms) and the fast repeated spirals (10 x 25 ms). The response post-photostimulation was not significantly different (independent t-test for 1 x 250 ms vs. 10 x 25 ms: t=-1.06, p=0.293).

Firstly, we wanted to determine if the opsin was more effectively activated by repeatedly spiralling the neuron with fast spirals (ten 25 ms spirals) or spiralling with a single slow (250 ms) spiral. For each neuron, we spiralled the beam over the cell at the 0 µm z-position while simultaneously imaging the GCaMP6s response at 920 nm. We spiralled the same neurons with fast and slow spirals, alternating which stimulation was tested first for each neuron. There was no significant difference in calcium response between slow spiralling and multiple fast spirals (**Figure 2c**). However, we did this test at 6 mW average power, which was potentially already a saturating power. We decided to use slow spirals (1 x 250 ms) for the experiments as this produced a slightly lower response (independent t-test for 1 x 250 ms vs. 10 x 25 ms: t=-1.06, p=0.293).

We adjusted the average power of the photostimulation beam in 1 mW steps from 2 to 5 mW, as we had previously noticed that 6 mW was resulting in very large, possibly saturating responses (>3 ΔF/F; data not shown). Photostimulation of each neuron was repeated for three trials at each power level. The results showed that 4 and 5 mW average power of the OPSF achieved similar responses, potentially indicating that the neurons had reached a maximal response at this power (**Figure 2b**). We chose 3 mW average power for all further photostimulation experiments as the responses at 2 mW were too similar to control regions (∼100 µm from the targeted neuron; see Methods).

In summary, we chose an average power that produced sub-maximal responses at the central photostimulation plane (z = 0 µm) to ensure that we could accurately measure changes in neuronal GCaMP6s response at different z-positions.

### Measuring the effect of OPSF size on neuronal photostimulation precision

To test whether the size of the OPSF affects the precision with which neurons can be targeted *in vivo*, we stimulated single neurons at different z-positions with the two sizes of OPSF. We again used mice expressing GCaMP6s and C1V1-Kv2.1 in neurons of the somatosensory cortex (**Figure 2a**). We varied the z-position of the photostimulation OPSF while simultaneously imaging the GCaMP response from the targeted neuron at z = 0 µm, repeating three trials at each photostimulation z-position. To account for any effects of opsin desensitisation due to repeated exposure, the photostimulation z-position was either sequentially moved from +60 µm to -60 µm (‘standard’ direction) or from -60 µm to +60 µm (‘reverse’ direction; **Figure 3a**). Neurons were only tested for a single OPSF size and a single direction (standard or reverse), this was varied for each new cell to sample all of the combinations. The resultant response from each neuron was recorded and the normalised change in fluorescence (ΔF/F) was measured post-stimulation (n=8 cells for each direction for small OPSF and n=7 cells for reverse, n=8 cells for standard for the large OPSF). The neuronal responses were highest at positions around z = 0 µm and dropped to undetectable levels at the extreme z-positions (**Figure 3b, c**), implying a complete estimation of the entire response curve.

**Figure 3:**
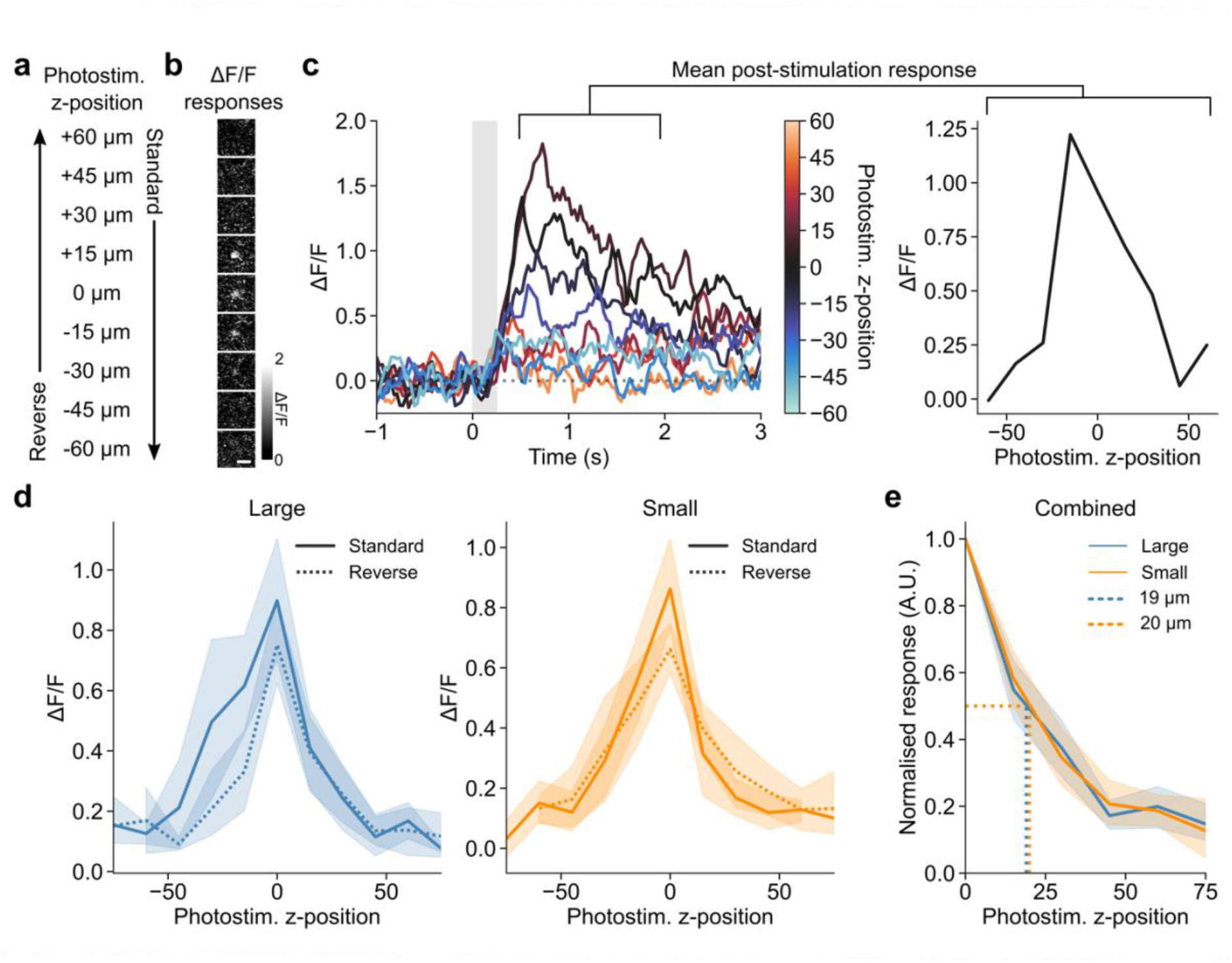
The accuracy of neuronal photostimulation in mouse somatosensory cortex layer 2 is not affected by variations in commonly used OPSF sizes. **(a)** Schematic of the different directions of photostimulation z-travel. The photostimulation OPSF was stepped sequentially through the z-dimension from top-to-bottom (+60 to -60 µm in 15 µm steps; ‘standard’) or bottom-to-top (‘reverse’) by changing the hologram produced by the SLM. Three trials were repeated at each z-position before moving to the next one. **(b)** Example mean ΔF/F images of post-photostimulation responses (mean of 1.5 seconds after stimulation end) for a single targeted neuron that was photostimulated at each z-position while being imaged at a fixed imaging plane (z = 0 µm). *Scale bar* = 20 µm. **(c)** *Left*, ΔF/F traces from a single targeted neuron showing responses at each photostimulation z-position. *Right*, the ΔF/F values in the period 1.5 seconds post-photostimulation were averaged to produce a neuronal response curve for axial photostimulation precision. **(d)** *Left*, curves showing mean post-photostimulation responses +/- 95% C.I. at each photostimulation z-position for the large OPSF, for the two different directions (different cells for each direction; reverse, n=7 cells; standard, n=8 cells). The standard and reverse conditions were not statistically different (independent t-test for standard versus reverse at each z-position: p>0.05). *Middle*, curves from the small OPSF (reverse, n=8 cells; standard, n=8 cells). *Right*, mean normalised photostimulation response +/- 95% C.I. versus photostimulation z-position, a summary of the previous four curves for the large and small OPSFs. The half-width at half-maximum is indicated (*dashed line*), and not different for the two curves (independent t-test for large versus small OPSFs et each z-position: p>0.05).

We compared the response curves from the standard and reverse directions of both OPSF sizes. Neurons that failed to respond with >0.5 ΔF/F in at least one z-position were removed, so that only responsive neurons were included. Additionally, due to the variation in the axial position of the imaging plane relative to the centre of the targeted neuron soma, we aligned individual response curves by their maximum response (**Figure 3d**). The maximum offset of a curve was one z-position (15 µm), which is about the size of a cell body and fits with the expected degree of uncertainty about axial position when imaging neuronal cell bodies. That is, the identification of a cell’s axial position in the imaging plane was difficult to determine at a low zoom (required to avoid imaging-induced photostimulation; see Methods), and so the z = 0 µm plane may have been at the very top or bottom of a cell. The standard and reverse directions were not significantly different from each other for either OPSF size (independent t-test for standard vs. reverse at each z-position: p>0.05), so they were combined to improve statistical power. Similarly, the two halves of the curve (above and below the cell) were combined, however we do note that the shape of the curve above the neuron was steeper for both OPSF sizes. The full-width at half-maximum (FWHM) of the photostimulation response curve for the large and small OPSFs was 38 and 40 µm respectively (independent t-test for large vs. small OPSFs et each z-position: p>0.05; **Figure 3e**).

Our results show that the PPSF does not depend on the axial extent of the OPSF for the OPSFs we measured and the biological system we utilised (pyramidal cells in layer 2 of mouse neocortex). To further understand this result we modelled the contribution of cell size to the axial extent of the PPSF (**Figure 4**). We used the width of a square pulse to model the cell diameter and convolved it with the axial extent of the large and small OPSFs to give the PPSF (**Figure 4b**). The results show that as the cell diameter increases beyond the size of the OPSF axial FWHM, the PPSF axial FWHM begins to depend solely on the cell diameter (**Figure 4c**). Beyond a certain size (∼30 µm), both OPSF sizes converge on the same PPSF, breaking the dependence of PPSF on OPSF.

**Figure 4:**
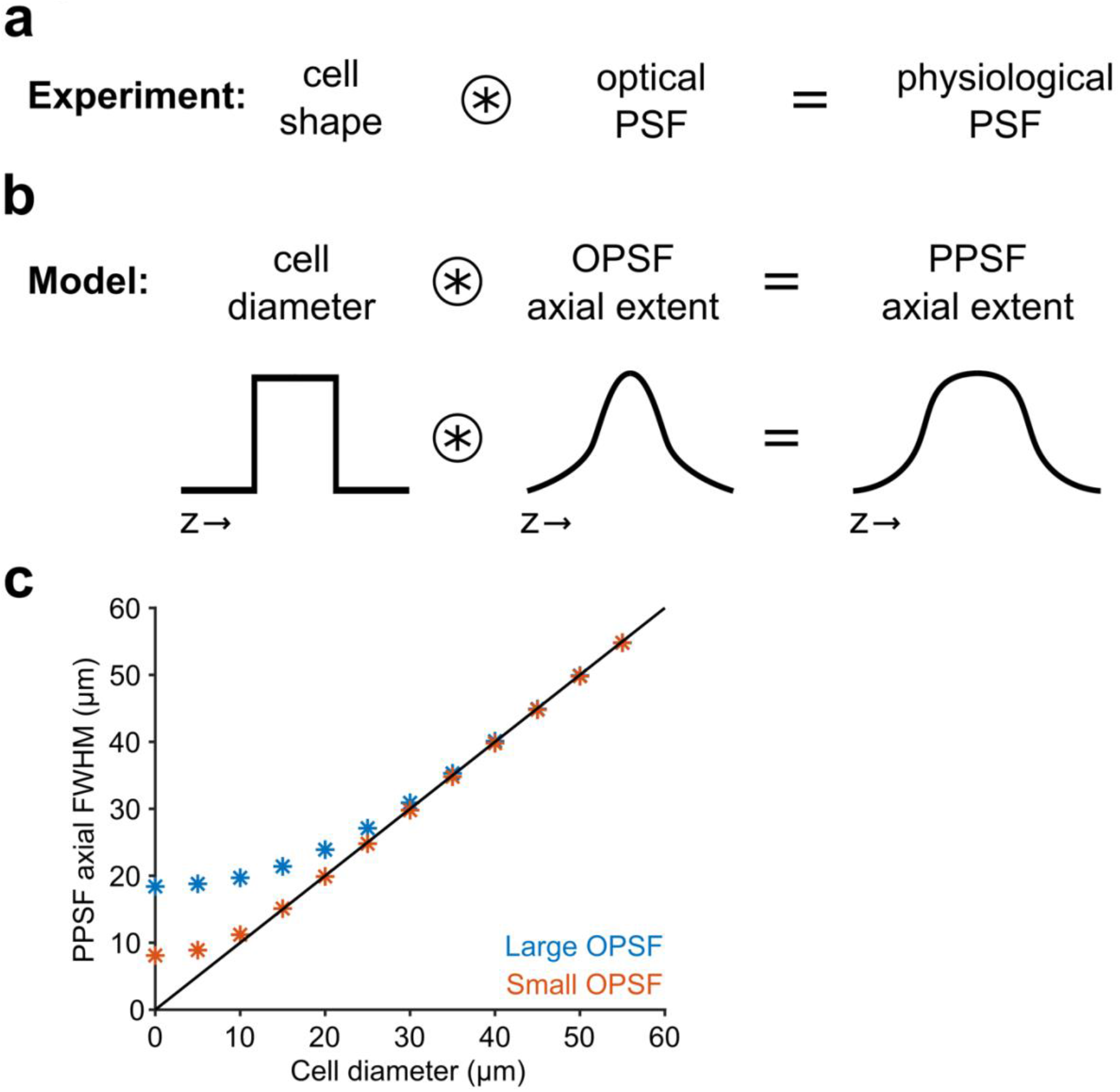
Modelling the physiological PSF for different cell shapes. **(a)** The PPSF is a result of the convolution of the cell shape and the optical PSF shape. **(b)** The cell diameter was used to model the cell shape as a sphere and the axial full width at half maximum (FWHM) from the fluorescence signal of the OPSF was used to model the contribution of the OPSF. The cell diameter was modelled as a square wave and the axial FWHM as a Gaussian. **(c)** A plot of the relationship between PPSF and cell diameter for the large (*blue stars*) and small (*orange stars*) OPSF. The line of equality is also plotted (*solid black line*).

## Discussion

We found that the axial FWHM of the PPSF during two-photon optogenetic excitation was not significantly affected when using OPSFs differing by more than a factor of two, from 8 µm to 19 µm. It can be challenging and expensive to achieve a tight OPSF using an SLM; the results presented here indicate that putting effort into improving OPSF beyond a certain limit does not increase the accuracy of the PPSF. This was further confirmed by modelling the PPSF as a convolution of the axial extent of the OPSF and the cell diameter, showing that the PPSF converges for large cells when using the different OPSFs from this study.

The laser power often employed for two-photon excitation of opsins may be “saturating” in two related ways. There may be greater photon flux per excitation volume than required to: excite all the opsin molecules in that volume (i.e. saturating light dosage per excitation volume), or excite all the opsin molecules in the somatic and nearby membranes (e.g. saturating activation of the opsin population per cell). Some kind of saturation seems likely given that the two-photon absorption cross-section of channelrhodopsin-2 is high, which is likely the case for other opsins as well. These facts further imply that there is effective out-of-focus excitation, as shown for isolated cells^3^. Out-of-focus excitation further sacrifices resolution beyond the minimum set by the physics alone, which would be the OPSF convolved with the cell, resulting in resolution dominated by the size of the cell (assuming OPSF is much less than cell size).

There are some limitations to our work. These results may be partly specific to the cell type, opsin, and light targeting methodology used here, although some principles described further in this Discussion can be generalised. Different types of cells, e.g. neurons from different species and/or different brain areas with different shapes and sizes, will require different parameters. Variation in the expression of opsin and GCaMP also likely add variability to our measurements, and could make it challenging to compare our study to other studies using different expression strategies.

Further, regarding the modelling of cell size versus PPSF, the contribution of cell shape in our model could be adjusted to include the dendritic compartments, which would have a large contribution to PPSF when using opsin restricted nearby, but not solely, to the cell body. Additionally, the opsin is found only in the membrane, and not across the entire cell volume, therefore a better model of the cell shape could encompass the varying amount of opsin across the cell. A larger surface area at the middle of an approximately spherical cell will contain more opsin molecules than at the poles.

Future work developing stronger opsins may help if they achieve their “strength” via increasing the conductance per protein, although this may also increase imaging-induced photostimulation in all-optical settings. However, simply increasing expression level globally would not help because then the increased photocurrent from somatic activation will scale in the same way that increased photocurrent due to out-of-focus excitation of perisomatic regions increases. Truly somatic restriction instead of just perisomatic enhancement would be most beneficial. Crucially, further optimisation of the spatial resolution at the optical level will not be beneficial (although see the limitations above).

In summary, the way forward to achieve optimal resolution may be to highly express (within healthy limits) high conductance opsin proteins in the somatic membrane alone. High expression and high conductance are desired to achieve sufficient depolarisation to reach action potential threshold without needing to activate all the opsin molecules and therefore avoid saturation in order to prevent the concomitant out-of-focus excitation. Care must also be taken to avoid imaging-induced photostimulation. Then, tailor the light to approximate a delta function (infinite at the center, zero everywhere else) in the axial direction as much as possible, at least such that the distribution of light is confined to a slab much thinner than the diameter of the target cells.

## Methods

### Animals, viruses and surgery

All animal experimentation was carried out with approval from the UK Home Office and University of Oxford Animal Welfare and Ethical Review Board. Three wildtype (C57Bl/6) mice and three mice transgenically labelled with GCaMP6s in excitatory neurons (B6;DBA-Tg(tetO-GCaMP6s)2Niell/J x CamK2a-tTa(AI94)) were used in this study.

Briefly, a cranial window, consisting of two #1 thickness coverglasses (3 mm and 4 mm diameter) adhered to one another using UV-curing optical adhesive, was implanted over the somatosensory cortex (-1.9 mm anterior and +3.8 mm lateral from Bregma) of each mouse during anaesthetised recovery surgery. During surgery, ∼800 nl of C1V1-Kv2.1 virus (AAV2/9-CaMKIIa-C1V1-t/t-kv2.1-mScarlet; ∼2 x 10^12 GC/ml diluted 1:5) was injected at 300 µm deep below the pial surface. In the case of wildtype mice, GCaMP6s virus (AAV2/1-syn-GCaMP6s-WPRE-SV40; ∼2.5 x 10^13 GC/ml diluted 1:10 in sterile phosphate-buffered saline) was also injected at the same time. The animals were allowed to recover for at least 3 weeks before imaging, which also allowed enough time for the virus to express in neurons.

### Microscopes and lasers

The microscope used in this study was a Scientifica HoloStim3D (from Scientifica, UK), equipped with a fixed wavelength 920 nm laser for imaging (Axon 920 from Coherent Inc. or FemtoFiber Ultra 920 from Toptica Photonics) and a 1030 nm fixed wavelength laser for photostimulation (Satsuma HP^2^ from Amplitude Laser), each with their own dedicated scan path. The HoloStim3D was equipped with a spatial light modulator (SLM; 1920 x 1152 calibrated for 1064 nm from Meadowlark Optics), which allowed for deflection of the photostimulation beam axially to the imaging plane without affecting the z-position of the imaging plane in the sample. ScanImage acquisition software (MBF Bioscience, Williston, VT), written in MATLAB, was used to control all imaging and photostimulation parameters. The microscope was equipped with a 16x magnification objective (Nikon 16x, 0.8 NA, N16XLWD-PF from Thorlabs), which was used throughout all calibrations, measurements and experiments.

### Varying and measuring optical point spread function

The effective numerical aperture of the photostimulation beam path was altered by cropping the 1920 x 1152 phase mask loaded onto the SLM. The undeflected zeroth order beam at the centre of the photostimulation field of view was blocked so that a single spot hologram could be measured from the first order deflections. The point spread function (PSF) of the photostimulation beam was estimated by imaging 0.2 µm beads (TetraSpeck Microspheres T7280 from ThermoFisher Scientific) that were dried on to a microscope slide using ethanol. Images for PSF estimation were acquired at 16x magnification, 256 x 256 pixels, 0.1 µm pixel size, 1 µm between slices in the axial dimension and 5 mW average power. Individual beads that were bright, and did not get damaged by the imaging process were cropped and chosen for PSF estimation. The resultant stacks were registered using the FIJI^33^ plugin StackReg^34^ to remove any tilt in the PSF. The MetroloJ^35^ plugin for ImageJ was used to fit a Gaussian curve to each individual bead in the axial and lateral dimensions. The curve was manually inspected for a good fit between the model and the data. At least 3 beads were imaged for each PSF to ensure results were consistent.

### Photostimulation

To determine whether fast versus slow spiralling was better, fast spiralling was done using 25 x 10 ms spirals, whereas slow spiralling used 1 x 250 ms spiral. For the comparison, each neuron was measured with both fast and slow spiralling, in a random order. Slow spiralling was used for all experiments. 6 mW average laser power was used for photostimulation in these experiments.

Subsequently, to find the best photostimulation laser power, different neurons were targeted for each power of 2, 3, 4 and 5 mW average power, no neuron was repeated with multiple laser powers.

For all further experiments, the 1030 nm photostimulation laser was used at 3 mW average power as measured by a sensor (S130C from Thorlabs) at the objective. The power was measured separately for each axial position and for each effective numerical aperture adjustment. The repetition rate of laser pulses was set to 2 MHz, with ∼350 fs pulse width and ∼4300 W peak power, the energy per pulse was ∼1.5 µJ. The SLM was kept at a temperature of ∼25 degree celsius.

Each axial position for the photostimulation beam was controlled by adjusting the diffraction from the SLM to focus the hologram deeper or shallower relative to the imaging plane. The first order deflection was ∼100 µm lateral to the zeroth order, which was blocked at a conjugate plane with a piece of metal to reflect/absorb the beam. The exact positions were calibrated and measured by burning a spot into a plastic slide (Autofluorescent Plastic Slides from Chroma) at each depth. The spots ranged axially from -60 µm to +60 µm relative to the imaging plane in steps of 15 µm, 9 steps in total.

Using separate galvanometers from the imaging path, the single spot hologram was spiralled with 5 concentric revolutions across the target neuron with a diameter of 10 µm at 3 mW average power. Repeats were carried out every 10 seconds, with 3 repeats per neuron per axial position. Different neurons were chosen for each change in optical resolution. The interval between each axial position was at least 30 seconds to allow GCaMP and opsin responses to recover. Stimulations of each neuron alternated between the ‘standard’ direction from -60 to +60 µm or the ‘reverse’ direction from +60 to -60 µm.

### Imaging

Responses to photostimulation were imaged at 30 Hz using resonant scanning and a wavelength of 920 nm. Images consisted of 512 x 512 pixels with a pixel size of ∼1.25 µm. Two separate detectors (multi-alkali PMTs) were used to collect emission from C1V1-Kv2.1-mScarlet and GCaMP6s. Group delay dispersion (GDD) was optimised for the imaging laser using a fluorescent sample to determine the best two-photon excitation of the sample. A large imaging FOV of 650 µm square was chosen to reduce the time that the imaging beam was scanning each neuron, reducing imaging-induced photoactivation of C1V1. An average imaging laser power of 20 mW was chosen to further reduce the effects of imaging-induced photoactivation, this power level was determined by scanning at different imaging powers and measuring the change in GCaMP fluorescence over a 30-second period. A power of 20 mW resulted in no immediate or slow ramp up of activity across the duration of imaging (data not shown).

During *in vivo* experiments, animals were head-fixed while awake and images were acquired at 150 µm deep in the somatosensory cortex (estimated layer 2). The cranial window was flattened by tipping/tilting the animal holder so that the window was perpendicular to the objective lens, resulting in less distortion of the images from refraction.

All imaging and photostimulation was synchronised using the voltage input/output recording software, PackIO^36^. A voltage trigger was used to begin imaging and photostimulation was manually triggered from ScanImage. Both the frame clock and galvanometer output were recorded to detect spiral initiation times relative to imaging frames.

### Data analysis

The voltage recordings of the frame clock and photostimulation trial onsets were used to find trial start times. Mean raw fluorescence trials were calculated across the three repeats for each neuron. Trials from different axial photostimulation positions were aligned to each other to ensure there was no drift between different acquisitions in the same field of view (FOV). Normalised change in fluorescence was calculated using ΔF/F = (F - F_0_)/F_0_ where F = the raw fluorescence, F_0_ = mean F in the baseline period (1.5 s before the stimulation onset). A freehand region of interest was drawn around the stimulated neuron and the mean ΔF/F of all of the pixels was taken. Control ROIs were drawn ∼100 µm away from the targeted neuron with the same shape as the neuronal ROI and analysed in the same way. The response for each trial was taken as the mean ΔF/F of the period post-stimulation (1.5 s after stimulation ended). Any neuron that responded with less than 0.5 ΔF/F across all axial positions of photostimulation was removed as it suggested that the photostimulation was unsuccessful. Overall, 15/31 neurons were included for the large optical point spread function and 17/38 neurons for the small one.

Due to the uncertainty of the exact position of the neurons in the axial dimension, the photostimulation precision curves were offset so that the maximum response was re-labelled as 0 µm. No curve was shifted by more than 15 µm (one axial position). The photostimulation precision for the different optical resolutions was measured as the full width half maximum (FWHM; the width of a horizontal line intercepting the signal at half the prominence of the peak of the curve) using the *find_peaks* and *peak_widths* functions as part of SciPy’s signal processing module for Python. Standard (-60 to +60 µm) and reverse (+60 to -60 µm) photostimulation response curves were combined by flipping the reverse curves so that the positions matched the standard condition.

All error bars are +/- 95% confidence interval, unless otherwise stated.

### Modelling

MATLAB was used to generate mock data for an axial OPSF from the experimental results. For each OPSF, a standard deviation was calculated from the FWHM value and a Gaussian distribution was generated from random numbers using that standard deviation. The data was normalised to the maximum values and a moving average filter of 5 was applied. Cell sizes were modelled as binary square waves, where the y-axis was 1 across the entire cell diameter on the x-axis. Each OPSF was convolved with the square wave of each cell size using the *convolve* function. The code and processed data is freely available to repeat these measurements, found here: https://github.com/Packer-Lab/OPSF_vs_PPSF.

## Author contributions

R.M.L and A.M.P. designed the study. R.M.L. conducted experiments. R.M.L. and B.P. performed analysis with advice from A.M.P.. R.M.L., B.P. and A.M.P. wrote the paper.

## Acknowledgements

We thank Kelly Sakaki, Simon Butt and Anna Hoerder-Suabedissen for their useful discussion. We also thank Scientifica, Toptica Photonics and Coherent Inc. for technical support with the microscopes and lasers. This work was supported by funding from the Wellcome Trust (204651/Z/16/Z) and the European Research Council (ERC) under the European Union’s Horizon 2020 research and innovation program (grant agreement No 852765).

## References

1. Oron, D., Papagiakoumou, E., Anselmi, F. & Emiliani, V. Two-photon optogenetics. in Progress in Brain Research vol. 196 119–143 (Elsevier, 2012).

2. Andrasfalvy, B. K., Zemelman, B. V., Tang, J. & Vaziri, A. Two-photon single-cell optogenetic control of neuronal activity by sculpted light. Proc. Natl. Acad. Sci. 107, 11981–11986 (2010).

3. Rickgauer, J. P. & Tank, D. W. Two-photon excitation of channelrhodopsin-2 at saturation. Proc. Natl. Acad. Sci. 106, 15025–15030 (2009).

4. Papagiakoumou, E., de Sars, V., Oron, D. & Emiliani, V. Patterned two-photon illumination by spatiotemporal shaping of ultrashort pulses. Opt. Express 16, 22039 (2008).

5. Packer, A. M. et al. Two-photon optogenetics of dendritic spines and neural circuits. Nat. Methods 9, 1202–1205 (2012).

6. Prakash, R. et al. Two-photon optogenetic toolbox for fast inhibition, excitation and bistable modulation. Nat. Methods 9, 1171–1179 (2012).

7. Rickgauer, J. P., Deisseroth, K. & Tank, D. W. Simultaneous cellular-resolution optical perturbation and imaging of place cell firing fields. Nat. Neurosci. 17, 1816–1824 (2014).

8. Packer, A. M., Russell, L. E., Dalgleish, H. W. P. & Häusser, M. Simultaneous all-optical manipulation and recording of neural circuit activity with cellular resolution in vivo. Nat. Methods 12, 140–146 (2015).

9. Emiliani, V., Cohen, A. E., Deisseroth, K. & Hausser, M. All-Optical Interrogation of Neural Circuits. J. Neurosci. 35, 13917–13926 (2015).

10. Pégard, N. C. et al. Three-dimensional scanless holographic optogenetics with temporal focusing (3D-SHOT). Nat. Commun. 8, 1228 (2017).

11. Oron, D., Tal, E. & Silberberg, Y. Scanningless depth-resolved microscopy. Opt. Express 13, 1468 (2005).

12. Zhu, G., van Howe, J., Durst, M., Zipfel, W. & Xu, C. Simultaneous spatial and temporal focusing of femtosecond pulses. 7 (2005).

13. Mardinly, A. R. et al. Precise multimodal optical control of neural ensemble activity. Nat. Neurosci. 21, 881–893 (2018).

14. Marshel, J. H. et al. Cortical layer–specific critical dynamics triggering perception. Science 365, eaaw5202 (2019).

15. Sridharan, S., et al. High performance microbial opsins for spatially and temporally precise perturbations of large neuronal networks. http://biorxiv.org/lookup/doi/10.1101/2021.04.01.438134<x> (2021) doi:10.1101/2021.04.01.438134.

16. Lim, S. T., Antonucci, D. E., Scannevin, R. H. & Trimmer, J. S. A Novel Targeting Signal for Proximal Clustering of the Kv2.1 K+ Channel in Hippocampal Neurons. Neuron 25, 385–397 (2000).

17. Wu, C., Ivanova, E., Zhang, Y. & Pan, Z.-H. rAAV-Mediated Subcellular Targeting of Optogenetic Tools in Retinal Ganglion Cells In Vivo. PLOS ONE 8, e66332 (2013).

18. Baker, C. A., Elyada, Y. M., Parra, A. & Bolton, M. M. Cellular resolution circuit mapping with temporal-focused excitation of soma-targeted channelrhodopsin. eLife 5, e14193 (2016).

19. Zhang, Z., Russell, L. E., Packer, A. M., Gauld, O. M. & Häusser, M. Closed-loop all-optical interrogation of neural circuits in vivo. Nat. Methods 15, 1037–1040 (2018).

20. Bounds, H. A. et al. Ultra-precise all-optical manipulation of neural circuits with multifunctional Cre-dependent transgenic mice. 2021.10.05.463223 Preprint at https://doi.org/10.1101/2021.10.05.463223 (2022).

21. Oldenburg, I. A., et al. The logic of recurrent circuits in the primary visual cortex. http://biorxiv.org/lookup/doi/10.1101/2022.09.20.508739<x> (2022) doi:10.1101/2022.09.20.508739.

22. Papagiakoumou, E. et al. Scanless two-photon excitation of channelrhodopsin-2. Nat. Methods 7, 848–854 (2010).

23. Carrillo-Reid, L., Yang, W., Bando, Y., Peterka, D. S. & Yuste, R. Imprinting and recalling cortical ensembles. Science 353, 691–694 (2016).

24. dal Maschio, M., Donovan, J. C., Helmbrecht, T. O. & Baier, H. Linking Neurons to Network Function and Behavior by Two-Photon Holographic Optogenetics and Volumetric Imaging. Neuron 94, 774–789.e5 (2017).

25. Shemesh, O. A. et al. Temporally precise single-cell-resolution optogenetics. Nat. Neurosci. 20, 1796–1806 (2017).

26. Forli, A. et al. Two-Photon Bidirectional Control and Imaging of Neuronal Excitability with High Spatial Resolution In Vivo. Cell Rep. 22, 3087–3098 (2018).

27. Forli, A., Pisoni, M., Printz, Y., Yizhar, O. & Fellin, T. Optogenetic strategies for high-efficiency all-optical interrogation using blue-light-sensitive opsins. eLife 10, e63359 (2021).

28. Yang, W., Carrillo-Reid, L., Bando, Y., Peterka, D. S. & Yuste, R. Simultaneous two-photon imaging and two-photon optogenetics of cortical circuits in three dimensions. eLife 7, e32671 (2018).

29. Gill, J. V. et al. Precise Holographic Manipulation of Olfactory Circuits Reveals Coding Features Determining Perceptual Detection. Neuron S089662732030578X (2020) doi:10.1016/j.neuron.2020.07.034.

30. McRaven, C., et al. High-throughput cellular-resolution synaptic connectivity mapping in vivo with concurrent two-photon optogenetics and volumetric Ca ^2+^ imaging. http://biorxiv.org/lookup/doi/10.1101/2020.02.21.959650<x> (2020) doi:10.1101/2020.02.21.959650.

31. Dalgleish, H. W. et al. How many neurons are sufficient for perception of cortical activity? eLife 9, e58889 (2020).

32. Daie, K., Svoboda, K. & Druckmann, S. Targeted photostimulation uncovers circuit motifs supporting short-term memory. http://biorxiv.org/lookup/doi/10.1101/623785<x> (2019) doi:10.1101/623785.

34. Schindelin, J., et al. Fiji: an open-source platform for biological-image analysis. Nat. Methods 9, 676–682 (2012).

34. Thevenaz, P., Ruttimann, U. E. & Unser, M. A pyramid approach to subpixel registration based on intensity. IEEE Trans. Image Process. 7, 27–41 (1998).

35. Matthews, C. & Cordelieres, F. P. MetroloJ: an ImageJ plugin to help monitor microscopes’ health. ImageJ User Dev. Conf. Proc. (2010).

36. Watson, B. O., Yuste, R. & Packer, A. M. PackIO and EphysViewer: software tools for acquisition and analysis of neuroscience data. bioRxiv (2016) doi:10.1101/054080.

